# Positive serial dependence in ratings of food images for appeal and calories

**DOI:** 10.1101/2024.02.07.579264

**Authors:** David Alais, Thomas Carlson

**Affiliations:** School of Psychology, The University of Sydney, Australia

## Abstract

Food is fundamental to survival and our brains are highly attuned to rapidly process food stimuli. Neural signals show foods can be discriminated as edible or inedible as early as 85 ms after stimulus onset^1^, distinguished as processed or unprocessed beginning at 130 ms^2^ and as high or low density from 165 ms^3^. Recent evidence revealed specialised processing of food stimuli in the ventral visual pathway^4-6^, an area that underlies perception of faces and other important objects. For many visual objects, present perception can be biased towards recent perceptual history (known as serial dependence^7,8^). We examined serial dependence for food in two large samples (n>300) that rated sequences of food images for either ‘appeal’ or ‘calories’. Calorie ratings by males and females agreed closely but appeal ratings were higher in males. High calorie ratings were associated with high appeal, especially in males. Serial analyses testing if current trial ratings were influenced by the previous one showed both appeal and calorie ratings exhibited clear positive dependences (i.e., a high preceding rating increased current trial ratings). The serial effect for appeal was roughly twice that for calories and males showed a greater serial effect than females for both ratings. Serial amplitude was larger in those who reported a longer elapsed time since they last ate and was larger in the BMI>25 group compared to BMI<25. These findings square with recently found food selectively in visual temporal cortex, reveal a new mechanism influencing food decision-making and suggest a new sensory-level component that could complement cognitive strategies in diet intervention.

## Results

Underlying the brain’s rapid analysis of food stimuli is a large, distributed network composed of numerous regions, including decision-related areas in prefrontal cortex and visual areas in occipital cortex. Among prefrontal areas, appetizing food stimuli activate several regions including orbitofrontal cortex^9-11^, medial prefrontal cortex^9,12^ and anterior cingulate^13^, with food saliency being linked particularly to orbitofrontal cortex^14^ and anticipation of pleasure from the food associated with lateral orbitofrontal cortex^15^. There is also sensory-driven food-related activity in visual cortex, including the lateral occipital complex and fusiform gyrus^3,16-18^, two visual object processing areas^10^. The strongest findings come from very recent studies analysing fMRI responses to large image sets which found highly selective food specialisation in the ventral visual pathway. Khosla et al.^5^ found distributed activation from food images that was not due to low-level image properties (i.e., colour, shape, texture). Jain et al.^4^ found food-specific activity adjacent to the fusiform face area. Another study found food images drove color-biased ventral visual areas and predicted voxel responses^6^.

The involvement of prefrontal cortex highlights the role of choice and decision-making in food behaviours. What to eat, when and how much are critical decisions, whether for survival (e.g., meeting caloric needs when resources are scarce) or in affluent Western societies where a surfeit of food choices leads us to make around 200 food decisions daily^19^. Following recent findings of robust food activity in ventral visual areas^4-6^, visual perceptual factors likely also influence food choice. This is relevant as recent perceptual history can bias current perceptual decisions so that they are not independent but biased towards recent input (known as ‘serial dependence’^20,21^). Many visual stimuli elicit this serial bias, from basic attributes (orientation, motion, spatial frequency, numerosity^22-25^) to more complex visual objects such as faces^26-28^, visual scenes^29^ and even artworks^30^. In particular, faces – another image category with high ecological significance processed in the fusiform gyrus – show strong serial biases for decisions about attractiveness, sex, identity and emotion^26,27,31-34^. Given that serial dependence involves high-level (decisional) and low-level (sensory) factors^35-38^, and that food is processed in frontal decisional and visual sensory regions, food images should also evoke serial biases. Here we examine how rating food for appeal or calorie content affects subsequent food ratings. We find clear evidence of positive serial dependences for both calories and appeal, with current ratings biased towards previous ratings.

### Food image ratings

Figure 2 shows ratings data for calories (2a) and appeal (2c). Data points are mean ratings from male (blue) and female (red) participants for all 150 images. Calorie ratings show close agreement between male and female participants and the ratings span nearly the full range of the scale. There was no significant difference between male and female ratings for calories (males: M = 46.612; females: M = 46.719; t_298_ = .036, p = .971. See Fig 2b). Appeal ratings showed a slightly different pattern. There was a significant difference between males and females (males: M = 54.630; females: M = 50.689; t_298_= 2.525, p = .012. See Fig 2d) and the range of ratings was reduced and clustered around the centre of the scale (compare appeal and calories standard deviations: Fig. 2b,d). Figure 2e,f plots the mean calorie and appeal ratings given for each of the 150 images by male (2e) and female (2f) participants. There was a significant correlation (higher calorie ratings associated with higher appeal) in both groups, with males (r =.559, p < .0001) showing a stronger association than females (r = .323, p = .0001).

**Figure 1.**
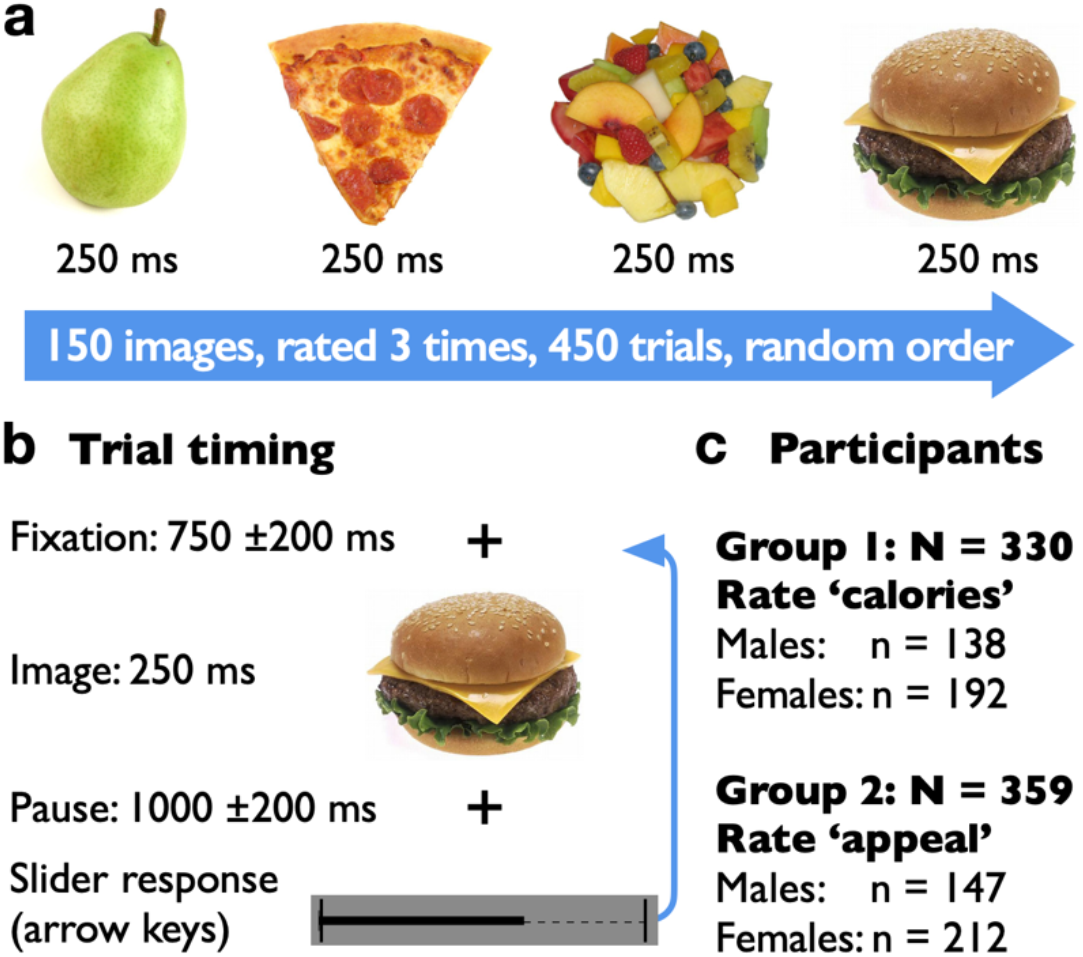
Stimuli and methods. (A) 150 food images were selected from the ‘transformed’ and ‘natural food’ categories of the FRIDa image set (https://foodcast.sissa.it/neuroscience). The set of 150 images was presented in a random order and then presented again two more times in new random orders (3 ratings per image). (B) Each trial began with a blank screen averaging 750 ms (±200 ms random jiQer), then a food image (250 ms) then a blank screen (1000 ms ±200), then a response slider that participants controlled with keyboard arrow keys to indicate their rating of the food (recorded with a mouse click). (C) Two groups were recruited from Prolific so that two dependent variables could be measured. Group 1 (n = 330) rated the calorie value of the food; group 2 (n = 359) rated the appeal of the food.

**Figure 2.**
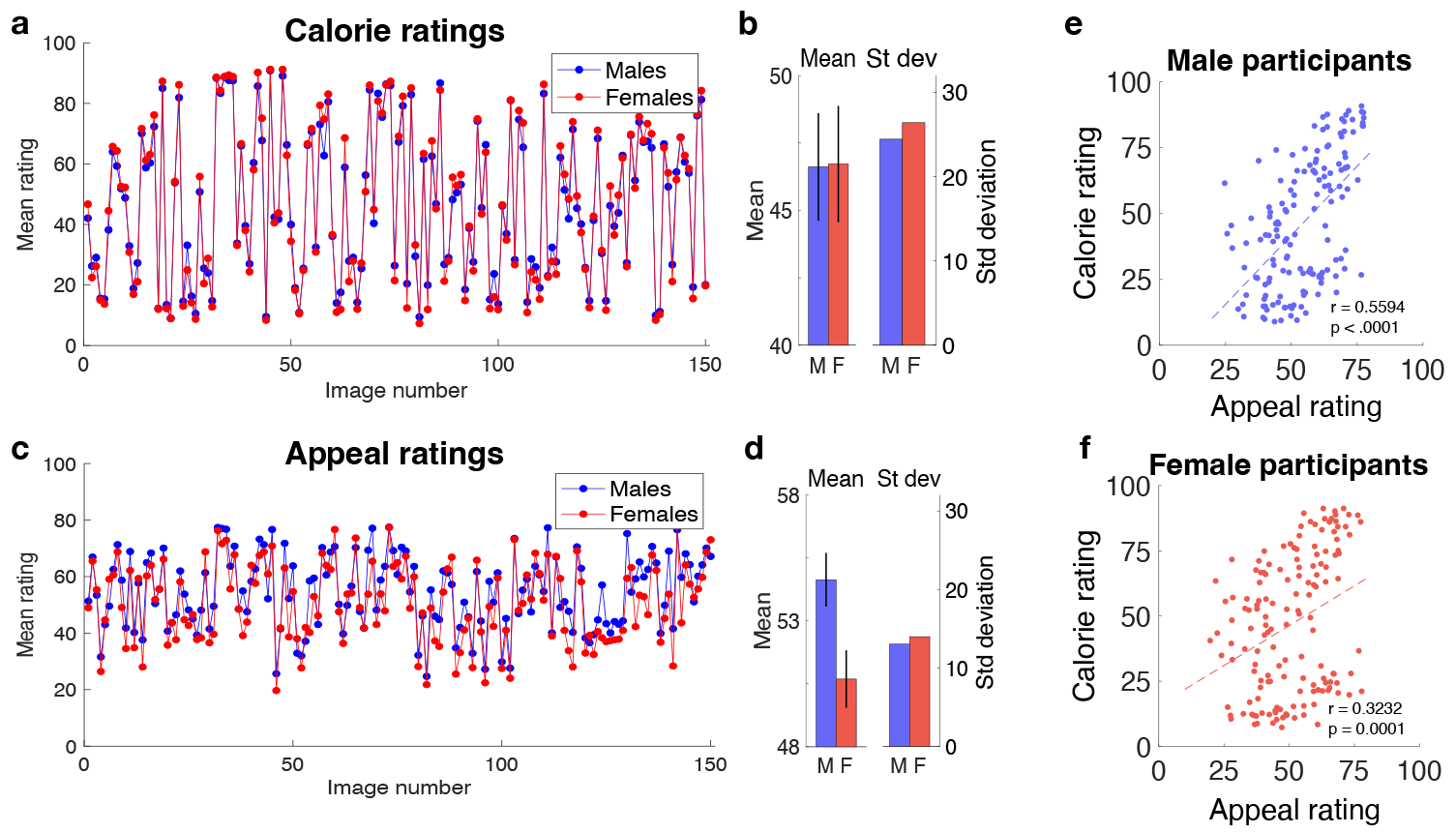
Mean ratings of 150 food images for calories and appeal. Group 1 (n=330: M = 138,F = 192) made calorie ratings (2a,b) and Group 2 (n=359: M = 147,F = 212) made appeal ratings (2c,d). Calorie ratings did not differ between males and females and were well spread over the scale range (see 2b). Ratings for food appeal did show a significant male/female difference (2d), with mean male appeal ratings being higher. Error bars in 2b,d are ±1 SEM. Mean calorie versus appeal ratings are shown in 2e,f. Each scatter plot contains 150 data points, being mean ratings for the 150 food images by male (2e) and female (2f) participants. Ratings are significantly positively correlated (higher calorie food rated as more appealing) for males and females.

### Serial dependence in food ratings

We next analysed the ratings data for serial effects to test whether successive responses were independent or whether instead a given trial’s response was influenced by the previous one. Figure 3 plots serial bias against relative difference. Relative difference (previous rating minus current rating) simply quantifies whether the previous rating was less than or greater than the current rating. Serial bias is effectively a deviation score quantifying how different the current image’s rating is from the mean of all ratings of that image. If all ratings are sequentially independent, then serial bias will be approximately zero for all values of relative difference. It is frequently observed, however, that perception on a current trial is biased *towards* the previous trial (a *positive* serial dependence). In the current experiment, this would manifest as a tendency to rate images higher when the previous rating was high, and lower when the previous rating was low.

**Figure 3.**
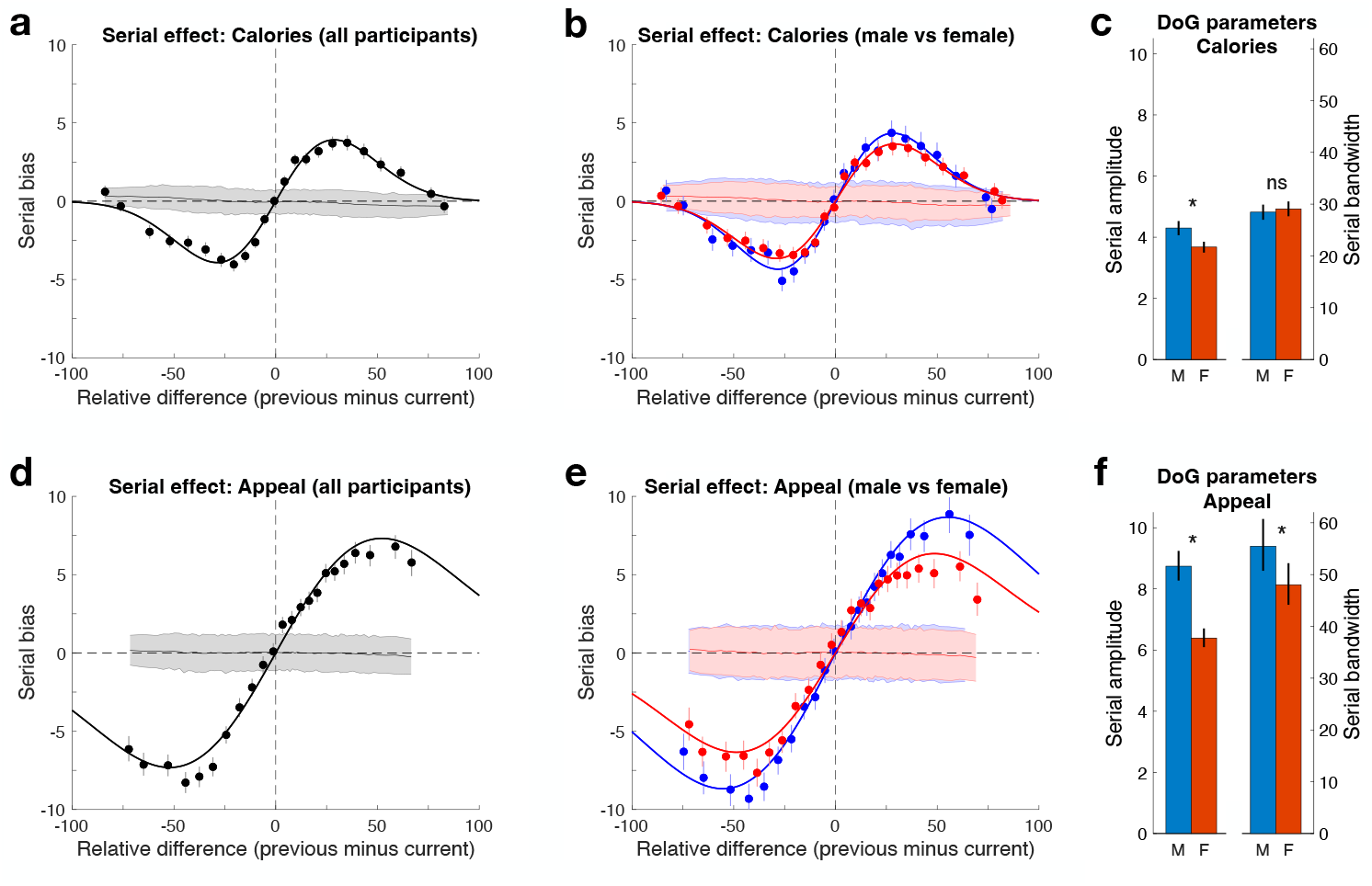
Food ratings for calories and appeal show a positive serial dependence. a) Calorie ratings deviate positively from their mean value when the previous food was rated highly (and negatively when the previous food was rated low). This aQraction to the previous food’s rating returns toward baseline for large differences between current and previous trial ratings and is well described by a Difference of Gaussian (DoG) model. The shaded band around zero bias shows the null distribution for serial bias produced by permuting the data 1000 times and refitting a DoG each time. Permuting trial order removes any sequential effects and the band shows the central 95% of the null distribution. b) Group calorie ratings from 3a split into male and female subgroups fitted with DoG functions. c) DoG parameters compared for male and female groups. Columns show the mean amplitude and bandwidth based on 1000 bootstraps of the data and a refitting of the DoG on each iteration. d,e,f) Appeal data: Lower panels contain the same analyses as the upper panels. The summary in 3f shows the serial effect is larger overall for appeal than for calories and is stronger for males in both amplitude and bandwidth. Error bars in all panels show 95% confidence intervals.

The upper panels of Figure 3 show the serial analysis for calorie ratings. The analysis for all participants (3a) shows a classic serial dependence effect: current trial data was biased towards higher values when the previous trial rating was high (upper right quadrant) and was biased towards lower values when the previous trial rating was low (bottom left quadrant). For large relative differences the effect dissipates and returns to baseline (typical of serial dependence effects^24,35,39,40^) and is well described by a difference-of-Gaussian (DoG) model (here, r^2^ = 0.990) with an amplitude of 3.918 and a bandwidth of 28.769. The gray shaded band shows the 95% confidence interval around the serial effect, calculated from 1000 iterations of permuting the trial order and repeating the serial analysis. This provides a null distribution because the permutation will remove any serially dependent effects and therefore indicates the effect expected by chance. Here it is clear our serial effect is both systematic (conforming to the expected DoG function) and far exceeds the 95% confidence limit.

Figure 3b shows the same serial analysis of the calorie ratings after splitting the group into male and female participants. For both subgroups, the serial effect conforms well to the DoG model (males: r^2^ = 0.988; females: r^2^ = 0.989) and again the shaded bands around zero serial bias show the 95% confidence interval based on permuting the data. The DoG parameters for males and females are compared in Figure 3c which plots amplitude and bandwidth for calorie ratings together with 95% confidence intervals (based on 1000 bootstraps of the data and refitting the DoG function each time). Males and females did not differ in the bandwidth parameter (the range of relative difference over which the serial effect occurs: Males = 28.307, females = 28.987, p = .829) but the amplitude of the serial effect differed significantly, with males showing a larger serial effect (males = 4.341, females = 3.656, p < .001).

The lower panels of Figure 3 show the serial analysis for appeal ratings. Figure 3d confirms the serial effect also occurs for rating food on appeal and indeed the effect is larger in amplitude and broader in range than was observed for calorie ratings (amplitude = 7.316, bandwidth = 52.123, r^2^ = 0.993). Figure 3e shows the serial effect for appeal ratings separately for male and female participants. The effect again conforms well to a DoG function (male: r^2^ = 0.997, female: r^2^ = 0.988) and far exceeds the 95% confidence interval around the null distribution (shaded bands in 3d,e). There is again a larger serial effect for males than for females (males = 8.671, females = 6.343, p < .001) and also a difference in bandwidth, with males showing a serial effect over a broader range (males = 55.306, females = 48.664, p = .016). Figure 3f summarises the amplitude and bandwidth parameters from the best-fitting DoG functions to male and female appeal ratings with 95% confidence intervals.

Finally, we analysed demographic and self-report ratings provided prior to the experiment.

Participants gave their age, height and weight (from which BMI was calculated), and whether they were currently dieting (yes/no), as well as answering the following questions on a 0-100 scale: How *hungry* are you now? How *thirsty* are you now? How *tired* are you now? They also indicated in hours: How long since your last meal? How long since you last ate? To test if any of these measures modulated the serial effect, we divided the sample into two subgroups using a median split (e.g., high hunger vs low hunger). The exceptions were dieting, which was split based on a binary yes/no response, and BMI where we compared BMI > 25 with BMI <25. We conducted serial analyses separately on the high and low subgroups for each variable and determined if the strength of the serial effect differed by subtracting the amplitude of the low group from the high group. Results are shown in Figure 4 and any value greater than zero indicates a stronger serial effect in the ‘high’ subgroup.

**Figure 4.**
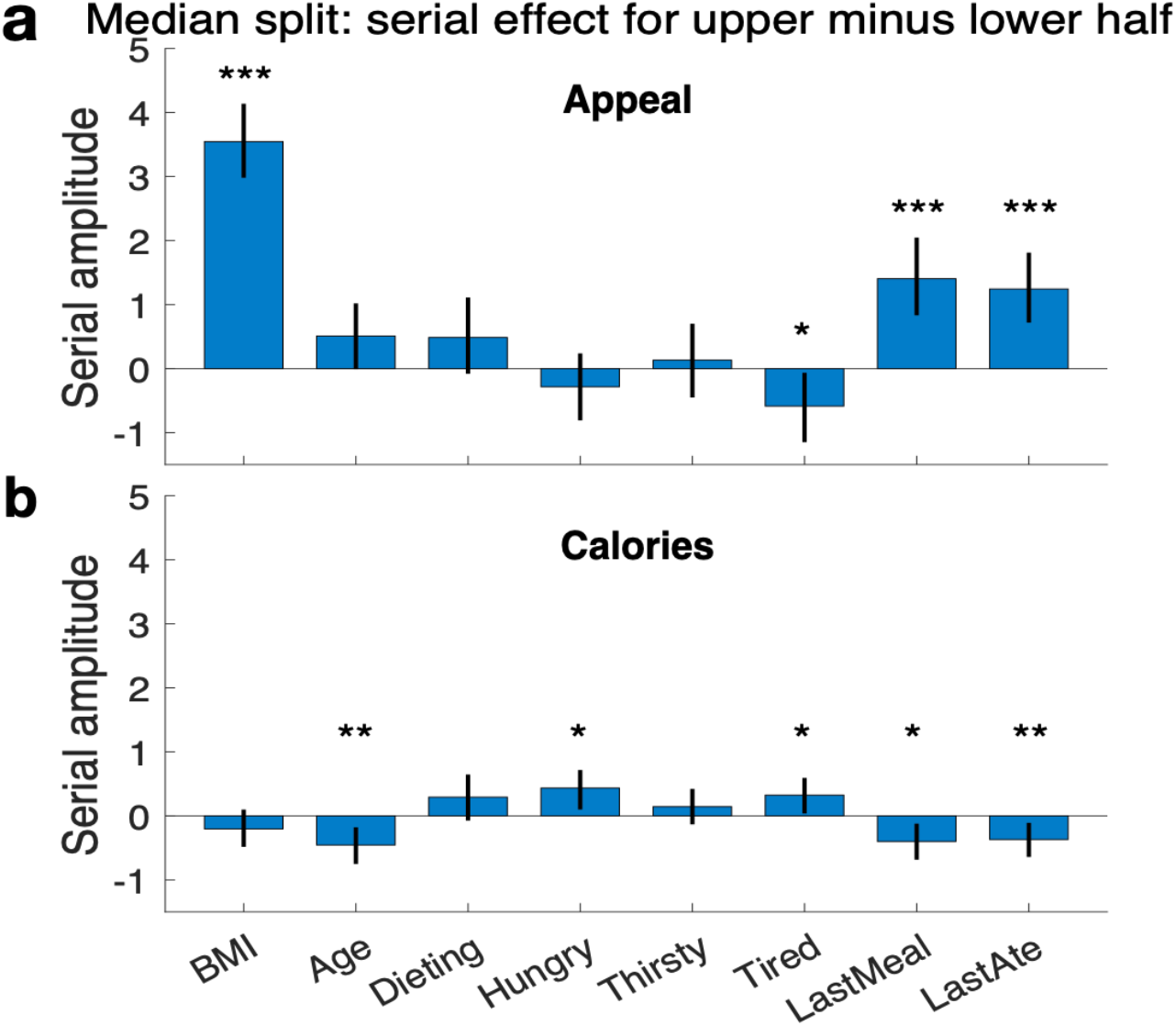
Results of dividing the sample using a median split. For each of the eight variables shown, the sample was divided into two subgroups and serial analyses were conducted for each group. Columns show the difference in the amplitude of the serial effect (high group minus low group), with values greater than zero indicating a stronger serial effect in the high group. A) For appeal ratings, there were strong and robust increases in serial amplitude for the high BMI group (BMI >25 vs BMI < 25) and in the groups that reported longer periods since they last ate or had a meal. B) Calorie ratings were much more stable between high and low groups for all eight variables, although the high statistical power of our large sample meant some small differences were significant. All error bars show 95% confidence intervals based on 1000 bootstraps.

## Discussion

This study was motivated by very recent findings showing that visual areas in human temporal cortex show a specialisation for food images^4-6^. We examined whether participants’ ratings of food images would exhibit a bias known as serial dependence, just as occurs with ratings of faces, another prominent visual specialisation in temporal cortex^41,42^ and for many other visual attributes^21^. Ratings of food images sequences for calories and appeal both showed significant positive serial dependences: rather than being sequentially independent, ratings tended to follow the rating given to the preceding image. A highly appealing preceding food image led to a current food image being rated higher than its mean appeal rating (and *vice versa*: a preceding low-appeal image made a current image less appealing). Rating food images for calories showed the same pattern of positive serial bias, although appeal ratings showed a stronger serial effect.

Serial dependence has been recently an active field in perceptual decision making. This attractive bias is an adaptive change in sensory information that makes current perception assimilate towards previously seen stimuli. It is widely reported in perceptual judgements about many stimuli from simple (e.g., such as orientation and spatial frequency^23,24^) to complex judgements about face perception (e.g., gender, attractiveness, identity^8^), scene perception^37^, and aesthetics^30^. Serial dependence also occurs at the level of global information where elements are combined into ensemble objects^29^ and may occur separately for individual objects in a multi-object scene^37,43^. It is also a real world-effect: when realistic movie clips are used, current perception of objects is biased by information presented up to 15 seconds earlier^44^ and current emotion is biased for up to 12 seconds^45^. These findings suggest that food serial dependence is likely to occur in real-world contexts containing multiple food options and future work could test this.

Food appeal showed a larger serial effect than food calorie ratings. Attention plays a role in boosting serial dependence effects^7,21^ and appealing foods capture attention, likely to a greater extent than an analytical food assessment such as rating calories. A related question is why serial dependence is greater in males. Perhaps males are more attentionally biased to food, driving a stronger serial effect. A recent review of the large literature on food attention bias notes few studies have reliable data on gender differences^46^ and some find no gender differences^47^. A systematic review found that food attentional bias is most prevalent in obese individuals^48^, which implies our gender differences are not BMI-related as median BMI scores were similar for males (24.8) and females (23.0). Follow-up work is needed to examine why food serial dependence differs depending on the rater’s gender and the food variable being rated.

Food decision-making is modulated by many factors^49-51^. We therefore collected data from participants on variables such as age, BMI, hunger, tiredness, etc. Comparing serial effects after a median split on each variable, the largest differences between high and low groups occurred for food appeal ratings (Fig. 4a), with the high BMI (>25) group showing greater serial dependence (nearly a factor of 2) than the low group and greater time since last eating or consuming a meal also driving stronger serial dependence. These findings fit with BMI exerting a direct influence on the brain’s food network^52,53^ and with elapsed time increasing hunger and thus food network activity^54^ and ghrelin levels (a hormone triggering hunger sensation and food seeking behaviours that increases neural response in orbitofrontal and visual cortex cortices^55^).

Given the high frequency of food decisions in our daily lives, serial dependence analyses are perfectly suited to uncovering drivers of food decision-making. We are surrounded by food options in affluent western countries, many highly processed with unhealthy combinations of calories, fat, sugar and salt. High ultra-processed food intake is linked with poor physical health (cardiovascular disease, cancer and overall mortality rate^56-58^) and mental health outcomes, with a recent meta-analysis finding greater odds of depressive and anxiety symptoms and increased risk of subsequent depression^59^. There is also evidence that such foods can induce eating addiction^60-62^. Understanding all aspects of food decision-making is therefore critical and our novel finding that food decisions exhibit positive serial dependence reveals a new component, one that adds a positive feedback loop making food look more appealing following a preceding appealing food, especially in overweight/obese individuals or those who have not eaten for several hours, and thus driving consumption. Conversely, food serial dependence makes a food less appealing when it follows a food rated low in appeal. Equivalently, the calorie rating given to a food could be boosted or attenuated by the preceding food. These findings could help guide clinical interventions, adding a sensory-based component to predominantly cognitive-based strategies designed to either reduce food intake (e.g., when treating obesity or compulsion to eat) or increase it (e.g., when treating bulimia or anorexia nervosa).

## Methods

### Participants

Experiment 1 measured calorie ratings and involved 330 participants; 138 males (42% of sample) and 192 females (58%). Experiment 2 measured appeal ratings and involved 359 participants; 147 males (41% of sample) and 212 females (59%). Recruitment was made online through Prolific and there were no exclusions during recruitment or data analysis.

### Stimuli

Food images were selected from the FRIDa image set (https://foodcast.sissa.it/neuroscience). We used 150 images drawn from the ‘transformed’ and ‘natural food’ categories (75 per category) that were selected to approximately evenly span the calorie range. These are all color images, cropped and presented on a white background, and were standardised to a size of 375 x 375 pixels and a resolution of 72 dpi. Stimulus presentation duration 250 ms.

### Procedure

The procedure was the same for Experiments 1 and 2. After instructions and practice trials to become familiar with the procedure participants were presented with 450 trials (a set of 150 food images presented three times with each set in a new random order). The trial sequence was: (i) a blank screen with a fixation cross for 750 ms (randomly varied in the range of ±200 ms), (ii) an image presentation for 250 ms, (iii) a blank fixation screen again for 1000 ms (randomly varied within ±200 ms), and (iv) a keyboard-driven ratings bar used to record the participant’s response. The ratings bar ranged from 0 – 100 with the slider initially set to 50 and needing to be adjusted before a response was accepted. In Experiment 1 (n=330) the task was to rate the foods for calories. In Experiment 2 (a different sample of n=359) the task was to rate the foods for appeal.

Demographic data were collected for every participant (age, gender, height, weight) and responses to the following: rate your hunger; rate your thirst; rate your tiredness; are you currently dieting?; how long since you last ate (hours)?; how long since your last meal (hours)?

### Design and data analysis

The aim of the study was to measure ratings for a large number of food images (n=150). Collecting data online meant that a large number of participants could be reached so that variability in demographic variables such as gender and BMI could be analysed. The use of a large set of food images has the potential limitation of relatively few ratings per image (3 ratings each) yet having a large sample means this is easily overcome by analysing data as single aggregate subject (a ‘super subject’ analysis) and using a bootstrapping procedure to obtain measures of variance and to calculate confidence intervals.

## STAR⋆Methods

### Key resources table

| REAGENT or RESOURCE | SOURCE | IDENTIFIER |
| --- | --- | --- |
| **Deposited data** |  |  |
| Psychophysical data | This paper | To be deposited on OSF (URL) |
| **Software and algorithms** |  |  |
| Online code (TC to upload) | This paper | To be deposited on OSF (URL) |
| MATLAB R2021a | This paper | RRID: SCR 001622 |
| Psychtoolbox-3 | This paper | RRID: SCR 002881 |

## Resource availability

### Lead contact

Further information and requests for resources should be directed to and will be fulfilled by the lead contact, David Alais.

## Materials availability

This study did not generate new unique reagents

## Data and code availability

Data reported in this study and the custom code for analyses are deposited at Open Science Framework (OSF URL). Custom code for experiment and visualization are available by request to the lead contact.

## Acknowledgments

This research was supported by a grant from the Australian Research Council (DP210101691) awarded to D.A..

## Author contributions

D.A. conceived of the study and D.A. and T.C. designed the research. T.C. implemented the experiment for online data collection. D.A. analysed and interpreted the data, prepared the figures and wrote the paper with feedback on the manuscript draft from T.C..

## Declaration of interests

The authors declare no competing interests.

## Supplemental information

None.

## Notes

### Competing Interest Statement

The authors have declared no competing interest.

